# Immunisation of ferrets and mice with recombinant SARS-CoV-2 spike protein formulated with Advax-SM adjuvant protects against COVID-19 infection

**DOI:** 10.1101/2021.07.03.451026

**Authors:** Lei Li, Yoshikazu Honda-Okubo, Ying Huang, Hyesun Jang, Michael A. Carlock, Jeremy Baldwin, Sakshi Piplani, Anne G. Bebin-Blackwell, David Forgacs, Kaori Sakamoto, Alberto Stella, Stuart Turville, Tim Chataway, Alex Colella, Jamie Triccas, Ted M Ross, Nikolai Petrovsky

**Affiliations:** Vaxine Pty Ltd., Bedford Park, Adelaide 5042, SA, Australia; College of Medicine and Public Health, Flinders University, Adelaide 5042, SA, Australia; Center for Vaccines and Immunology, University of Georgia, Athens, GA, USA; Department of Infectious Diseases, University of Georgia, Athens, GA, USA; Department of Pathology, University of Georgia, Athens, GA, USA; Centre for Virus Research, Westmead Millennium Institute, Westmead Hospital and University of Sydney, Sydney, 2145, NSW, Australia; School of Medical Sciences and Marie Bashir Institute, University of Sydney, Sydney, NSW 2006, Australia

## Abstract

The development of a safe and effective vaccine is a key requirement to overcoming the COVID-19 pandemic. Recombinant proteins represent the most reliable and safe vaccine approach but generally require a suitable adjuvant for robust and durable immunity. We used the SARS-CoV-2 genomic sequence and *in silico* structural modelling to design a recombinant spike protein vaccine (Covax-19™). A synthetic gene encoding the spike extracellular domain (ECD) was inserted into a baculovirus backbone to express the protein in insect cell cultures. The spike ECD was formulated with Advax-SM adjuvant and first tested for immunogenicity in C57BL/6 and BALB/c mice. The Advax-SM adjuvanted vaccine induced high titers of binding antibody against spike protein that were able to neutralise the original wildtype virus on which the vaccine was based as well as the variant B.1.1.7 lineage virus. The Covax-19 vaccine also induced potent spike-specific CD4+ and CD8+ memory T-cells with a dominant Th1 phenotype, and this was shown to be associated with cytotoxic T lymphocyte killing of spike labelled target cells *in vivo*. Ferrets immunised with Covax-19 vaccine intramuscularly twice 2 weeks apart made spike receptor binding domain (RBD) IgG and were protected against an intranasal challenge with SARS-CoV-2 virus 2 weeks after the second immunisation. Notably, ferrets that received two 25 or 50μg doses of Covax-19 vaccine had no detectable virus in their lungs or in nasal washes at day 3 post-challenge, suggesting the possibility that Covax-19 vaccine may in addition to protection against lung infection also have the potential to block virus transmission. This data supports advancement of Covax-19 vaccine into human clinical trials.

## Introduction

COVID-19 is caused by lung infection with severe acute respiratory syndrome coronavirus-2 (SARS-CoV-2)[1]. To date, there have been over 176 million reported COVID-19 cases in over 215 countries with greater than 3.8 million confirmed deaths [2]. SARS-CoV-2 causes a constellation of clinical outcomes from asymptomatic infection to respiratory failure and death [3]. Although public health strategies, such as social distancing, masks and quarantine, have helped control virus transmission, many countries are experiencing second and third waves of cases [4] including with virus variants of concern with increased virulence [5]. Global efforts are currently underway to develop COVID-19 vaccines [6] including live virus vectors, inactivated viruses, nucleic acids (DNA or RNA) and recombinant proteins [7]. Although several vaccines have received emergency-use authorisation, ongoing questions include likely duration of vaccine protection, long-term safety, potential for antibody-enhanced disease, activity against variant strains and immune correlates of vaccine protection [8–10].

Recombinant or inactivated protein vaccines are a safe and reliable approach but generally suffer from weak immunogenicity unless formulated with an appropriate adjuvant [11]. Adjuvants induce higher and more durable immune responses and can also be used to impart a relevant T helper bias to the immune effector response [12] and can overcome immune impairment seen with advancing age or chronic disease [13]. Advax-SM is a new combination adjuvant consisting of delta inulin polysaccharide particles (Advax™) formulated with the Toll-like receptor 9 (TLR9)-active oligonucleotide, CpG55.2. A similar adjuvant approach provided enhanced protection of recombinant spike protein vaccines against severe acute respiratory syndrome (SARS) and Middle East respiratory syndrome (MERS), coronaviruses [14, 15]. The Th1-bias imparted by the adjuvant also prevented the eosinophilic lung immunopathology otherwise seen after immunisation with SARS spike protein alone or with alum adjuvant [14]. Advax adjuvant has been shown to be safe and effective in human vaccines against seasonal and pandemic influenza [16, 17] and hepatitis B [18].

Advanced computer modelling techniques may be useful to accelerate pandemic vaccine design. This study describes how we used a range of approaches including computer modelling to characterise the SARS-CoV-2 spike protein from the genomic sequence, and then used a modelled 3-D structure to identify angiotensin converting enzyme 2 (ACE2) as the relevant human receptor. We then utilised our computer model to design a vaccine from the extracellular domain (ECD) of the SARS-CoV-2 spike protein, to test the hypothesis that this antigen when formulated with Advax-SM adjuvant would induce neutralising antibodies able to block the binding of the SARS-CoV-2 virus to ACE2 thereby preventing infection. While our study was in progress others confirmed that ACE2 was indeed the receptor for the spike protein and viral entry into host cells was further enhanced by priming of the spike protein by transmembrane protease serine 2 (TMPRSS2) [19]. Our subsequent results confirmed that our computationally-designed spike antigen when formulated with Advax-SM adjuvant induced antibodies against spike protein that were able to neutralise wildtype SARS-CoV-2 virus as well as pseudotyped lentivirus particles and cross neutralised the B.1.1.7 lineage virus. It induced memory CD4 and CD8 T cell responses with a Th1 phenotype and this translated into CTL killing of spike-labelled target cells *in vivo*. Notably, the Advax-SM adjuvanted vaccine when used in a ferret SARS-CoV-2 infection model protected against lung but also day 3 nasal virus replication, suggesting a potential ability to block virus transmission.

## Methods

### Vaccine Design

In mid-January 2020, we identified the putative spike protein from the SARS-CoV-2 genome sequence in NCBI (accession number: NC 045512) [20]. Given the homology of the spike proteins (76.4% sequence identity), SARS-CoV-1 was used as a template to model the SARS-CoV-2 spike protein. We performed a PSI-BLAST search against the Protein Data Bank (PDB) Database for 3D modelling template selection. Using the SARS-CoV-1 structure (PDB-ID 6ACC) [21] we performed structural homology modelling using Modeller9.23 (https://salilab.org/modeller/) to obtain a 3D structure of SARS-CoV-2 spike protein **(Figure 1A)**. The quality of the spike protein model was evaluated using GA341 and DOPE score, and the model was assessed using the SWISS-MODEL structure assessment server (https://swissmodel.expasy.org/assess). To help identify the putative cellular receptor for SARS-CoV-2, the crystal structure of human ACE2 (PDB-ID 3SCI) [22] was retrieved, and using HDOCK server, the spike protein was then docked against human ACE2 protein (http://hdock.phys.hust.edu.cn/) [23]. The docking poses were ranked using an energy-based scoring function and the docked structure analysed using UCSF Chimera. The high binding score predicted human ACE2 as the entry receptor for SARS-CoV-2 spike protein, confirming spike protein suitability for vaccine design [24]. The docked model was optimized using AMBER99SB-ILDN force field in Gromacs2020 (https://www.gromacs.org/). Molecular dynamic simulation (MDS) was carried out for 100 ns using a GPU-accelerated version of the program **(Supplementary video 1)**. The structural stability of the complex was monitored by the root-mean-square deviation (RMSD) value of the backbone atoms of the entire protein. The free energy of binding was calculated for simulated SARS-CoV-2 Spike and humanACE2 structure using g_MMPBSA. Finally, MDS was performed on the spike protein ECD vaccine construct to assess its ability to form a stable trimer despite the lack of the transmembrane and cytoplasmic domains. The insect cell codon-optimised expression cassette was cloned into pFASTBac1, and baculovirus was generated following standard Bac-to-Bac procedures. Recombinant baculovirus was expanded in Sf9 cells to P3 and then used for infection of High Five cells for protein expression. At 72h post infection, the culture supernatant was clarified by centrifugation, and then the recombinant spike protein ECD was purified by HisTrap Excel column using the AKTA chromatography system, concentrated by ultrafiltration and buffer changed to phosphate buffered saline (PBS) then terminally filter sterilised. The sequence of the recombinant spike protein (rSp) was confirmed by mass spectroscopy, SDS-PAGE gels and Western blots. Endotoxin was measured using a PyroGene™ Endotoxin Detection System (Cat. No. 50-658U, LONZA, Walkersville, MD, USA), and residual DNA content in the final vaccine product was also measured using a Quant-iT™ PicoGreen™ dsDNA Assay Kit (ThermoFisher, P7589) following manufacturer’s instructions.

**Figure 1:**
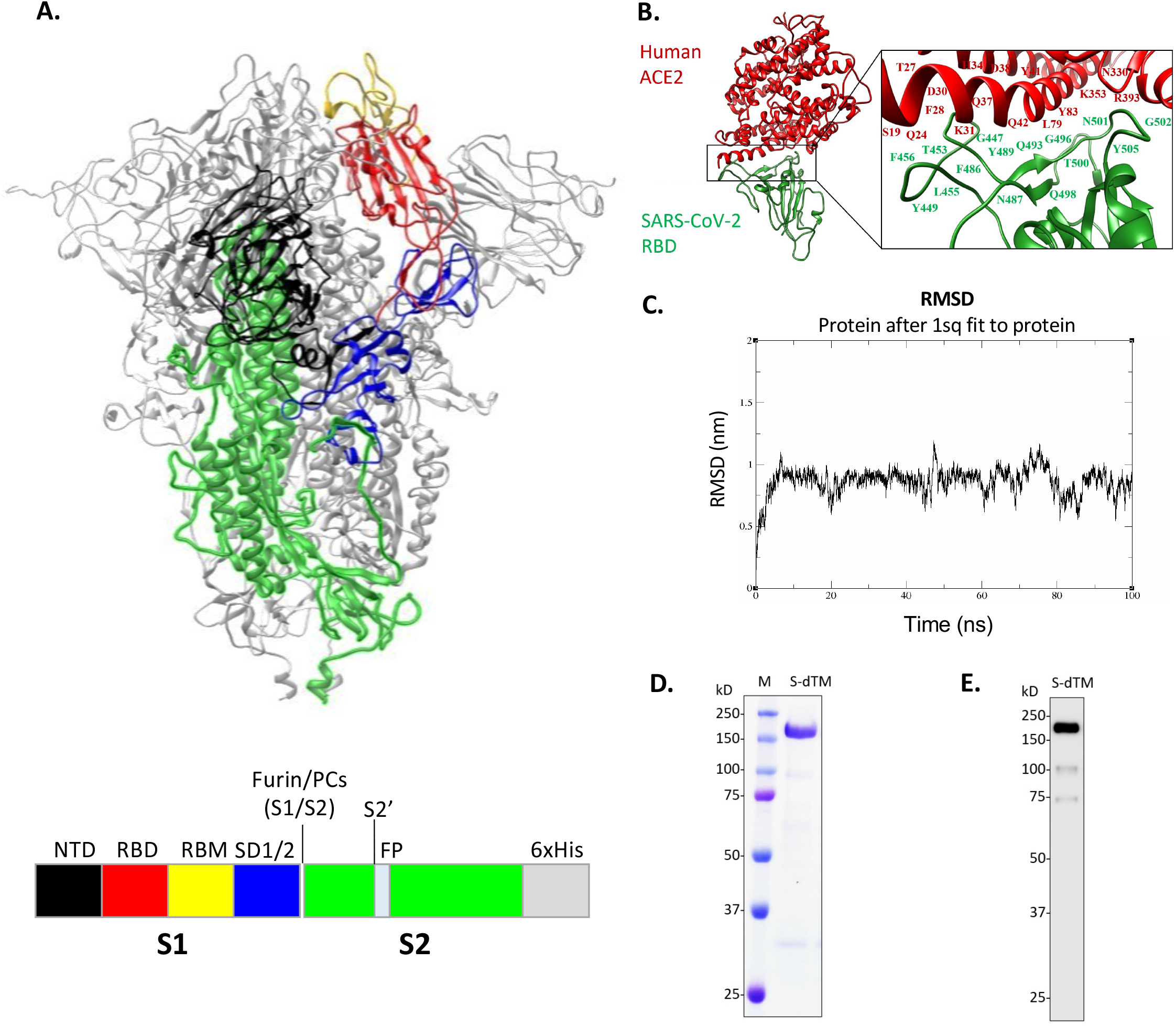
Modelling and expression of SARS-CoV-2 spike ectodomain (ECD) vaccine antigen. (A) Diagram of the ECD construct of SARS-CoV-2 spike (S) protein. Honeybee melittin signal (HBMss) peptide was used for efficient secretion expression. Transmembrane domain and intracellular domain were removed and replaced with a TEV cleavage site followed by 6x Histidines (6xHis). (B) SARS-CoV-2 receptor binding domain (RBD) (green) binding human Angiotensin-Converting Enzyme 2 (ACE2) (red). (C) Molecular dynamic simulation (MDS) run for 100 ns showed the structure was stable. (D) Representative Coomassie Brilliant Blue stained SDS-PAGE of purified S-dTM. (E) Purified S-dTM was confirmed by Western blot using anti-SARS-CoV-2 spike mouse polyclonal antibody.

### Mouse Immunisation Protocol

Female, BALB/c and C57BL/6 (BL6) mice (6-10 weeks old) were supplied by the central animal facility of Flinders University. Mice were immunised intramuscularly (i.m.) with 1 or 5 μg rSp alone or mixed with either 1 mg Advax-SM adjuvant or where indicated 50 μg Al(OH)_3_ (2% Alhydrogel, Croda Denmark) in the thigh muscle at weeks 0 and 2. Blood samples were collected by cheek vein bleeding at 2 weeks after each immunisation. Serum was separated by centrifugation and stored at −20°C prior to use. Advax-SM (Vaxine, Adelaide, Australia) was a sterile suspension of delta inulin microparticles at 50 mg/ml with CpG55.2 at 0.5 mg/ml.

### Antigen-specific ELISA for Murine Studies

Spike-specific antibodies were determined by ELISA. Briefly, 1 μg/ml rSp [corresponding to SARS-CoV-2 (Wuhan) reference sequence Q13 to P1209] or 2 μg/ml receptor-binding domain (RBD) [corresponding to SARS-CoV-2 (Wuhan) reference sequence R319 to F541] antigen in PBS were used to coat 96-well ELISA plates. After blocking, 100 μl of diluted serum samples were added followed by biotinylated anti-mouse IgG (Sigma-Aldrich), IgG1, IgG2a/c, IgG2b, IgG3 and IgM antibodies (all from Abcam) with horseradish peroxidase(HRP)-conjugated Streptavidin (BD Biosciences) for 1 hour (h). After washing, 100 μl of TMB substrate (KPL, SeraCare, Gaithersburg, MD, USA) was added and incubated for 10 min before the reaction was stopped with 100 μl 1M Phosphoric Acid (Sigma-Aldrich). The optical density was measured at 450 nm (OD_450_ nm) using a VersaMax plate reader and analysed using SoftMax Pro Software. Average OD_450_ nm values obtained from negative control wells were subtracted.

### High-throughput SARS-CoV-2 Live Virus Neutralisation Assay

A high-content microscopy approach was used to assess the ability of mice sera to inhibit SARS-CoV-2 viral infection and the resulting cytopathic effect in live permissive cells. Briefly, serum samples were diluted in cell culture medium (MEM-2% FCS) to create a 2-fold dilution series (e.g. 1:20 to 1:160). Each dilution was then mixed (in duplicate) with an equal volume of virus solution (B.1.319 or B.1.1.7 strains) at 8×10^3^ TCID_50_/ml (so that dilution series becomes 1:40-1:320), followed by 1 h incubation at 37°C. Meanwhile, freshly trypsinised VeroE6 cells were and plated in 384-well plates (Corning #CLS3985) at 5×10^3^ cells per well in 40 μL. After 1 h of virus-serum coincubation, 40 μL were added to the cell-plate for a final well volume of 80 μL. Plates were incubated for 72 h until readout (37°C, 5% CO_2_, >90% relative humidity), which occurred by staining cellular nuclei with NucBlue dye (Invitrogen, #R37605) and imaging the entire well’s area with a high-content fluorescence microscopy system (IN Cell Analyzer 2500HS, Cytiva Life Sciences). The number of cells per well was determined using InCarta image analysis software (Cytiva). The percentage of viral neutralisation for each well was calculated with the formula N = (D-(1-Q))x100/D, where “Q” is the well’s nuclear count divided by the average nuclear count of the untreated control wells (i.e. without virus or serum), and “D” equals 1 minus the average Q-value for the positive infection control wells (i.e. cells + virus, without serum). Therefore, the average nuclear counts for the infected and uninfected cell controls are defined as 0% and 100% neutralisation levels, respectively. The threshold for determining the neutralization endpoint titre of diluted serum samples mixed with virus was set to N≥50%.

### SARS-CoV-2 Spike Pseudotyped Neutralisation Assay

A non-replicative SARS-COV-2 Spike pseudotyped lentivirus-based platform was developed to evaluate neutralisation activity in infected/convalescent sera in a Biosafety Level 2 (BSL2) facility. The hACE2 open reading frame (Addgene# 1786) was cloned into a 3rd generation lentiviral expression vector pRRLSIN.cPPT.PGK-GFP.WPRE (Addgene# 122053), and clonal HEK 293T cells stably expressing ACE2 were generated by lentiviral transductions as described previously [25], followed by single cell sorting into 50% HEK 293T conditioned media (media conditioned from 50% confluent HEK 293T cultures). Lentiviral particles pseudotyped with SARS-COV2 Spike envelope were produced by co-transfecting HEK 293T cells with a GFP encoding 3rd generation lentiviral plasmid HRSIN-CSGW (a gift from Camille Frecha [26]), psPAX2 and plasmid expressing codon optimized but C-terminal truncated SARS COV2 S protein (pCG1-SARS-2-S Delta18 [27], herein Spike Delta18) courtesy of Professor Stefan Polhman using polyethylenimine as described previously [25]. Neutralisation activity of donor sera was measured using a single round transduction of ACE2-HEK 293T with Spike pseudotyped lentiviral particles. Briefly, virus particles were pre-incubated with serially diluted donor sera for 1 h at 37°C.

Virus-serum was then added onto ACE2-HEK 293T cells seeded at 2,500 cells per well in a 384-well tissue culture plate a day before. Following spinoculation at 1200xg for 1 h at 18°C, the cells were moved to 37°C for a further 72 h. Entry of pseudotyped particles was assessed by imaging GFP-positive cells and total cell numbers imaged through live nuclei counter staining using NucBlue (Invitrogen). Total cell counts and % GFP-positive cells were acquired using the InCell imaging platform followed by enumeration with InCarta high content image analysis software (Cytiva). Neutralisation was measured by reduction in % GFP expression relative to control group infected with the virus particles without any serum treatment.

### Murine T-cell response

BALB/c and BL6 mice were sacrificed, and individual spleens were collected one to two weeks after the last immunisation. Single-cell suspension in sterile PBS+3% FCS was prepared using a 70 μm easy strainer (Greiner Bio-One) with a 5 ml syringe plunger. Isolated spleen cells were pelleted and incubated in red blood cell (RBC) lysis buffer. For Cytometric Bead Array (CBA) assay, splenocytes were cultured at 5 x 10^5^ cells/well in 96-well plates with 3 μg/ml of rSp antigen [corresponding to SARS-CoV-2 (Wuhan) reference sequence Q13 to P1209] at 37°C and 5% CO_2_. Two days later, the supernatants were harvested and cytokine concentrations determined by mouse Th1/Th2/Th17 CBA (BD) and analysed by FCAP array Software (BD). In addition to CBA assay, enzyme-linked immune absorbent spot (ELISPOT) assay was performed using mouse Interlukin-2 (IL-2), Interlukin-4 (IL-4) or Interferon gamma (IFN-γ) ELISPOT set (BD PharMingen) or Interlukin-17 (IL-17) antibodies (BioLegend) according to the manufacturer’s instruction. Briefly, single-cell suspensions were prepared from spleens of mice and plated in Millipore MultiScreen-HA 96-well filter plates (Millipore) pre-coated with anti-mouse IL-2, IL-4, IL-17 or IFN-γ antibodies overnight at 4°C and blocked by RPMI-1640 containing 10% FBS. Cells were incubated for 48 h in the presence or absence of rSp protein at 37°C and 5% CO_2_. Wells were washed and incubated with biotinylated labelled anti-mouse IL-2, IL-4, IL-17 or IFN-γ antibody at room temperature (RT). After washing, wells were incubated with HRP-conjugated Streptavidin (BD Biosciences) for 1 h at RT. Wells were extensively washed again and developed with 3-amino-9-ethyl-carbazole (AEC) substrate set (BD Biosciences). After drying, spots were counted on an ImmunoSpot ELISPOT reader (CTL ImmunoSpot Reader, software version 5.1.36). Finally, a T-cell proliferation assay was performed by incubating collected mice spleen cells for 7 min at RT with 5 μM Carboxyfluorescein succinimidyl ester (CFSE) (Life Technologies), staining was quenched with FCS and splenocytes cultured at 10^6^ cells/well in 96-well plates for 5 days at 37°C in 5% CO_2_ with 3 μg/ml of rSp antigen. At the end of the incubation, the cells were stained with anti-mouse CD4-PerCP-Cy5.5 and anti-mouse CD8-APC (both from BD) and analysed on a FACSCanto II (BD). T-cell proliferation was expressed as the ratio of divided daughter cells to total T-cells, expressed as a percentage, by analogy to calculation of a stimulation index in thymidine proliferation assays.

### *In vivo* CTL assay

Functional CD8^+^ T cell response was determined by performing *in vivo* CTL assays, as described earlier [28]. Briefly, naïve syngeneic target spleen cells were left unpulsed or pulsed for 2 h in humidified CO_2_ incubator at 37°C with 5μM H-2K^b^-restricted Sp_539-546_ (VNFNFNGL) synthetic peptide [29] (DGpeptide, Hangzhou, China). Unpulsed (control) and peptide (antigen)-pulsed spleen cells were labelled with 0.5 μM CFSE (CFSE^low^) and 5 μM CFSE (CFSE^high^), respectively. Then, naïve syngeneic and immunised mice were adoptively transferred with 4×10^6^ cells of a 1:1 mix of control-to-antigen-pulsed target spleen cells. Eighteen (18) hours later, adoptive transfer recipient mice were euthanized, their splenocytes isolated and resuspended in PBS for acquisition on a BD FACSCanto-II instrument. To evaluate the percentage of antigen-specific target cell killing, the ratio of CFSE^high^/CFSE^low^ in survivors was compared to the ratio in transferred naive control mice.

### Ferret Immunisation Protocol

Fitch ferrets (*Mustela putorius furo*, spayed female, 6 to 12 months of age), were purchased from Triple F Farms (Sayre, PA, USA). Ferrets were pair-housed in stainless steel cages (Shor-Line, Kansas City, KS, USA) containing Sani-Chips laboratory animal bedding (P. J. Murphy Forest Products, Montville, NJ, USA) and provided Teklad Global Ferret Diet (Harlan Teklad, Madison, WI, USA) and fresh water ad libitum. Groups (n = 6) were vaccinated at day 0 and boosted at day 14 with Covax-19 vaccine (12.5, 25 or 50 μg) formulated with 15 mg Advax-SM adjuvant. Control ferrets received either saline only (n=3) or were immunised with influenza recombinant hemagglutinin vaccine (rH7, Protein Sciences, Meriden, USA) formulated with Advax-SM as a control. Blood was collected on days 0, 14 and 28 post-immunisation and day 10 post-challenge and stored at −20°C prior to use.

### Ferret Challenges

At day 28, all ferrets were infected intranasally with SARS-CoV-2 virus (1 × 10^5^ PFU) and were monitored daily during the infection for adverse events, including weight loss and elevated temperature for 10 days. At day 3 post infection, nasal swabs were collected from all animals, and three animals from each group, except for the Advax-SM only control group, were humanely euthanized, and lung tissue was collected. Three lobes from the right lung of each animal was formalin fixed for histopathology. Two lobes from the left side of lung from each animal were snap frozen and homogenised using 1 ml DMEM, and the supernatant was collected and kept frozen at −80° for viral titres.

### Ferret spike RBD-binding IgG ELISA

Immulon^®^ 4HBX plates (Thermo Fisher Scientific, Waltham, MA, USA) were coated with 100 ng/well of recombinant SARS-CoV-2 Spike protein RBD [corresponding to SARS-CoV-2 (Wuhan) reference sequence R319 to F541] in PBS overnight at 4°C in a humidified chamber. Plates were blocked with blocking buffer made up with 2% bovine serum albumin (BSA) Fraction V (Thermo Fisher Scientific, Waltham, MA, USA), 1% gelatin from bovine skin (Sigma-Aldrich, St. Louis, MO, USA) and 0.05% PBST (PBS with 0.05% Tween20) (Thermo Fisher Scientific, Waltham, MA, USA) at 37°C for 90 min. Serum samples from the ferrets were initially diluted 1:50 and further serially diluted 1:3 in blocking buffer to generate a 4-point binding curve (1:50, 1:150, 1:450, 1:1350). Subsequently, the plates were incubated overnight at 4°C in a humidified chamber. The following day, plates were washed 5 times with 0.05% PBST, and IgG antibodies were detected using horseradish peroxidase (HRP)-conjugated goat anti-ferret polyclonal IgG detection antibody (Abcam, Cambridge, UK) at a 1:4,000 dilution for a 90 min incubation at 37°C. Plates were washed 5 times with 0.05% PBST prior to colorimetric development with 100 μL of 0.1% 2,2’-azino-bis(3-ethylbenzothiazoline-6-sulphonic acid) (ABTS, Bioworld, Dublin, OH, USA) solution with 0.05% H_2_O_2_ for 18 min at 37°C. The reaction was terminated with 50 μL of 1% (w/v) sodium dodecyl sulfate (SDS, VWR International, Radnor, PA, USA). Colorimetric absorbance at 414 nm was measured using a PowerWaveXS plate reader (Biotek, Winooski, VT, USA). The dilution curve was plotted, and the area under the curve was calculated and multiplied by 1,000 to give standard units.

### Determination of Virus Titres in Ferret Nasal Washes

Nasal washes were titrated in quadruplicates in Vero E6 cells. Briefly, confluent VeroE6 cells were inoculated with 2-fold serial dilutions of sample in DMEM containing 2% FBS, supplemented with 1% penicillin-streptomycin (10,000 IU/ml). At 3 days post infection (dpi), virus positivity was assessed by reading out cytopathic effects. Infectious virus titres (TCID_50_/ml) were calculated from four replicates of each nasal wash using the Reed–Muench method.

### Haematoxylin & Eosin (H&E) and Immunohistochemistry Staining of Ferret Lungs

To assess the viral replication and pathological effect of infection, ferrets (n=3) were euthanised 3 days post infection. The right lung lobes were taken for viral plaques, the incision was tied with surgical suture, and the lung was inflated with 10 ml formalin. Lungs were removed and placed into formalin for 1 week prior to paraffin embedding. Ferret lungs were embedded into paraffin and were cut using a Lecia microtome. Transverse 5 μm sections were placed onto Apex superior adhesive glass slides (Leica biosystem Inc, IL, USA), which were coated for a positive charge, and were processed for H&E staining. Briefly, sections were deparaffinised in xylene and hydrated using different concentrations of ethanol (100%, 95%, 80% and 75%) for 2 min each. Deparaffinised and hydrated lung sections were stained with hematoxylin (Millipore sigma, MA, USA) for 8 min at RT, differentiated in 1% acid alcohol for 10 sec, and then counterstained with eosin (Millipore sigma, MA, USA) for 30 sec. Slides were then dehydrated with 95% and 100% ethanol, cleared by xylene, and mounted using Permount^®^ mounting media (Thermo Fisher scientific, MA, USA). Lung lesions were scored by a board-certified veterinary pathologist blinded to the study groups as follows: Alveolar (ALV) score: 1 = focal, 2 = multifocal, 3 = multifocal to coalescing, 4 = majority of section infiltrated by leukocytes; Perivascular cuffing (PVC) score: 1 = 1 layer of leukocytes surrounding blood vessel, 2 = 2-5 layers, 3 = 6 – 10 layers, 4 = greater than 10 cells thick; Interstitial Pneumonia (IP) score: 1 = alveolar septa thickened by 1 leukocyte, 2 = 2 leukocytes thick, 3 = 3 leukocytes, 4 = 4 leukocytes.

For lung immunohistochemistry, the deparaffinised and hydrated lung tissue sections were subjected to antigen retrieval by sub-boiling in 10 nM sodium citrate buffer at pH6 for 10 min and then incubated in 3% fresh-made hydrogen peroxide for 10 min to inactivate endogenous peroxidase at RT. The lung sections were blocked with 5% horse serum in PBS for 1 h at RT, incubated with SARS-CoV-2 Nucleoprotein polyclonal antibody at 1:500 dilution (Invitrogen, Carlsbad, CA, USA) overnight at 4°C, and then incubated with biotinylated goat-antibody Rabbit IgG H&L (Abcam, Waltham, MA, USA) at 1:1000 dilution for 1 h at RT. The avidin-biotin-peroxidase complex (VectStain Standard ABC kit, Vector Laboratories, Burlingham, CA, USA) was used to localise the biotinylated antibody, and DAB (Vector Laboratories, CA, USA) was utilised for colour development. Sections were then counterstained with hematoxylin, and then mounted using Permount^®^ mounting media (Thermo Fisher Scientific, Waltham, MA, USA). Images were obtained by Aperio digital slide scanner AT2 (Leica biosystem, Buffalo Grove, IL, USA).

### Statistical analysis

GraphPad Prism 8.3.1 for Windows was used for drawing graphs and statistical analysis (GraphPad Software, San Diego, CA, USA). The differences of antibody levels were evaluated by the Mann-Whitney test, and other differences between groups were evaluated by two-tailed Student’s t-test. ANOVAs with Dunnett’s test was used for weight loss with a statistical significance defined as a p-value of less than 0.05. Limit of detection for viral plaque titres was 50 pfu/ml for statistical analysis. Limit of detection for neutralisation is 1:10, but 1:5 was used for statistical analysis. Geometric mean titres were calculated for neutralisation assays. For all comparisons, p<0.05 was considered to represent a significant difference. In figures * = p < 0.05; ** = p < 0.01; and *** = p < 0.001. All error bars on the graphs represent standard mean error.

### Ethics statement

The mouse studies were performed at Flinders University, Australia. The protocol was approved by the Animal Welfare Committee of Flinders University and carried out in strict accordance with the Australian Code of Practice for the Care and Use of Animals for Scientific Purposes (2013). All efforts were made to minimise animal suffering. Animals were housed in cages provisioned with water and standard food and monitored daily for health and condition. After final monitoring, all of the surviving animals were humanely euthanised. Ferret studies were performed at The University of Georgia, United States. The University of Georgia Institutional Animal Care and Use Committee approved all ferret experiments, which were conducted in accordance with the National Research Council’s Guide for the Care and Use of Laboratory Animals, The Animal Welfare Act, and the CDC/NIH’s Biosafety in Microbiological and Biomedical Laboratories guide.

## RESULTS

### COVAX-19 Vaccine Design and Production

Based on the high binding score (−57.6 kcal/mol) seen from docking our 3D-model of spike protein to several putative receptors, we predicted ACE2 as the human entry receptor for SARS-CoV-2 **(Figure 1B)** [24]. This was soon confirmed by other groups using *in vitro* assays [19]. Based on this spike protein model, we sought to design a stable soluble secreted spike protein trimer for use as our vaccine immunogen. We designed a synthetic gene comprising the spike protein extracellular domain (ECD) together with N-terminal honeybee melittin signal sequence (HBMss) to ensure protein secretion and attached a hexa-histidine tag at the C-terminal end to assist with protein purification **(Figure 1A)**. Molecular dynamic simulation performed on the spike protein ECD vaccine construct confirmed its ability to form a stable trimer despite the lack of the transmembrane and cytoplasmic domains and the absence of any large trimerisation domain tag as used by others **(Figure 1C)**. The spike ECD gene construct was constituted into a baculovirus backbone, and the subsequent virus then used to transfect two insect cell lines (SF9 and Tni). While both cell lines successfully secreted the protein construct, higher protein expression was obtained in the Tni cells and these were used for subsequent production of a recombinant spike protein which was purified using a nickel affinity column and sterile filtration. The final protein product had a purity of ~ 90% by SDS-PAGE **(Figure 1D & E)** and was sterile with a low endotoxin and residual DNA content (data not shown).

### COVAX-19 vaccine induces spike protein RBD-binding and neutralising antibodies in mice

The serum anti-spike protein response of BL6 mice immunised with rSp alone was dominated by IgG1, a T helper 2 (Th2) isotype, whilst in Advax-SM adjuvanted groups the response was characterised by a switch to more IgG2b/c and IgG3 against spike **(Figure 2B)**. Overall, Advax-SM adjuvant was associated with a much higher anti-spike IgG2/IgG1 ratio consistent with a Th1-biased response **(Figure 2C)**. To provide an adjuvant comparison, we set up an additional study to compare the effects of the Advax-SM adjuvant to a traditional aluminium hydroxide adjuvant (Alhydrogel). As expected, the Alhydrogel adjuvant exacerbated the IgG1 (Th2) bias seen with rSp alone, thereby contrasting with the IgG2/3 (Th1) bias of the Advax-SM adjuvant. Most strikingly, mice immunised with Advax-SM adjuvanted rSp demonstrated high levels of *in vivo* cytotoxic T lymphocyte (CTL) killing of spike-labelled target cells, whereas mice immunised with rSp alone or formulated with Alhydrogel demonstrated minimal CTL activity against spike-labelled targets consistent with their Th2 immune bias **(Supplementary Figure 1)**.

**Figure 2:**
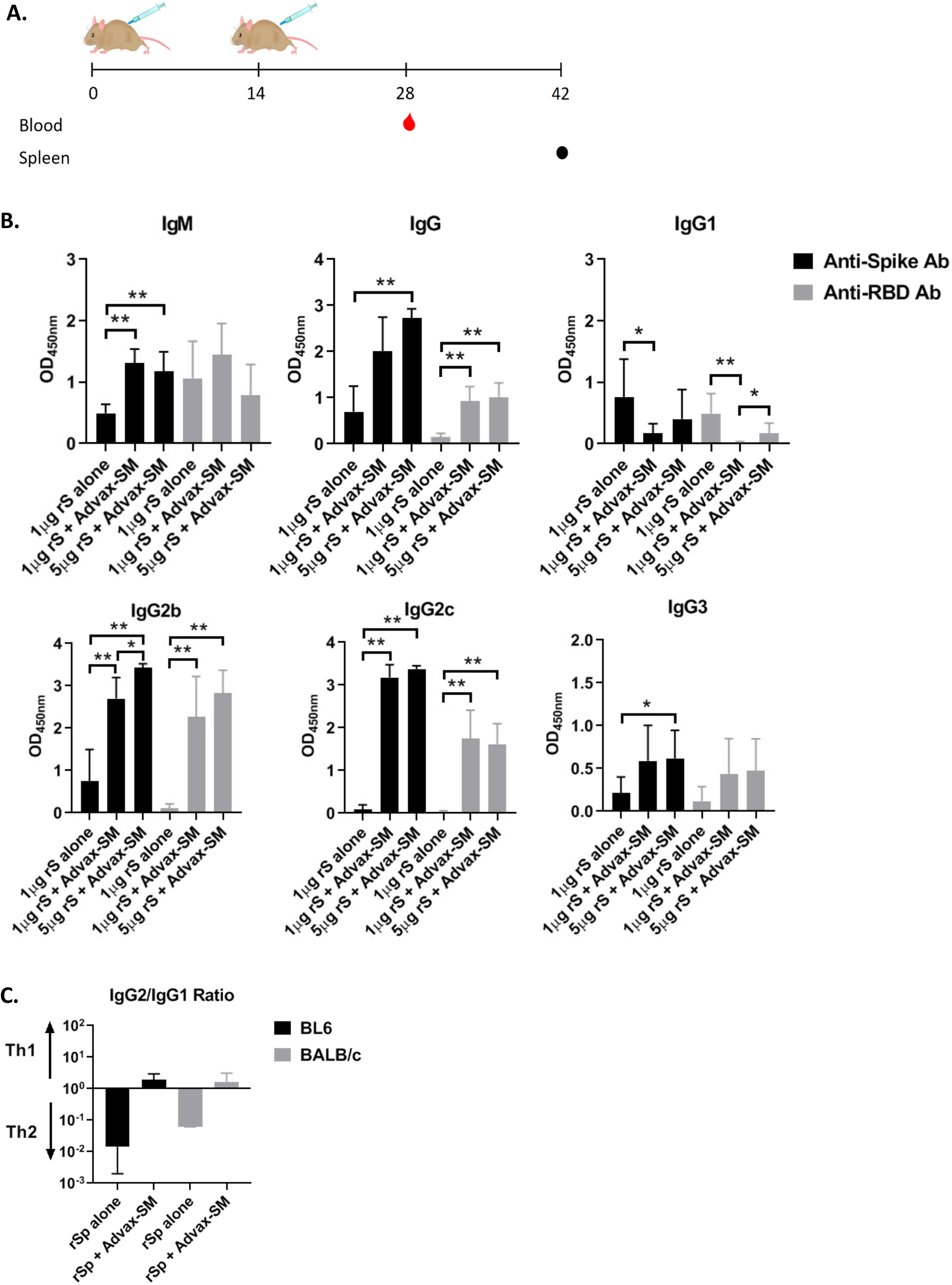
Advax-SM adjuvant enhances total Spike and RBD antibodies. (A) Schematic diagram of immunisation and sample collection. Female C57BL/6 and BALB/c mice were immunised i.m. twice at 2-week intervals with 1 μg rSp or 1 and 5 μg rSp with Advax-SM adjuvant. Blood samples were collected 2 weeks later and spleens 3-4 weeks after the last immunisation. (B) Shown are ELISA results (mean + SD) in BL6 mice. Statistical analysis by Mann-Whitney test. *; p < 0.05, **; p < 0.01, ns; not significant. (C) Overview of IgG2/IgG1 ratio in BL6 and BALB/c mice.

Spike RBD-binding antibodies have been reported to correlate with SARS-CoV-2 virus neutralisation [30]. There was a high correlation for each IgG subclass between the level of spike and RBD binding antibodies by ELISA, suggesting a significant proportion of spike antibodies induced by our rSp antigen were directed against the RBD region. RBD-binding IgG was almost undetectable in mice immunised with rSp alone, although these mice did exhibit some RBD-binding IgM. Notably, the Advax-SM adjuvant increased the spike IgG response but particularly favoured production of RBD-binding antibodies when expressed as a ratio of the total spike IgG response **(Figure 2C)**.

BALB/c mice, which have an overall Th2 bias, exclusively made IgG1 when immunised with rSp alone. Similar to what was seen in BL6 mice, in BALB/c mice Advax-SM adjuvanted rSp induced a switch from anti-spike IgG1 to IgG2b/c and IgG3 production The increased IgG2b/c and IgG3 induced by the Advax-SM adjuvant was equivalent to the reduction in IgG1. Interestingly, BALB/c mice had a low ratio of RBD to spike IgG, with IgM the dominant RBD-binding antibody rather than IgG. This contrasted with the high RBD to spike IgG ratio seen in the BL6 mice **(Supplementary Figure 2)**.

To determine whether the spike antibodies induced by our vaccine could neutralise virus infectivity, immune sera were tested in both a pseudotyped lentivirus assay (pseudovirus assay) and a SARS-CoV-2 neutralisation assay (live virus assay). BL6 mice immunised with rSp 1 μg with Advax-SM adjuvant showed significantly higher pseudovirus neutralisation titers (GMT 320) compared to BL6 mice immunised with an equivalent dose of spike protein alone (GMT 140) **(Figure 3A)** whereas BALB/c mice immunised with 1 μg rSp showed similar levels of pseudovirus neutralisation regardless of the presence of adjuvant.

**Figure 3:**
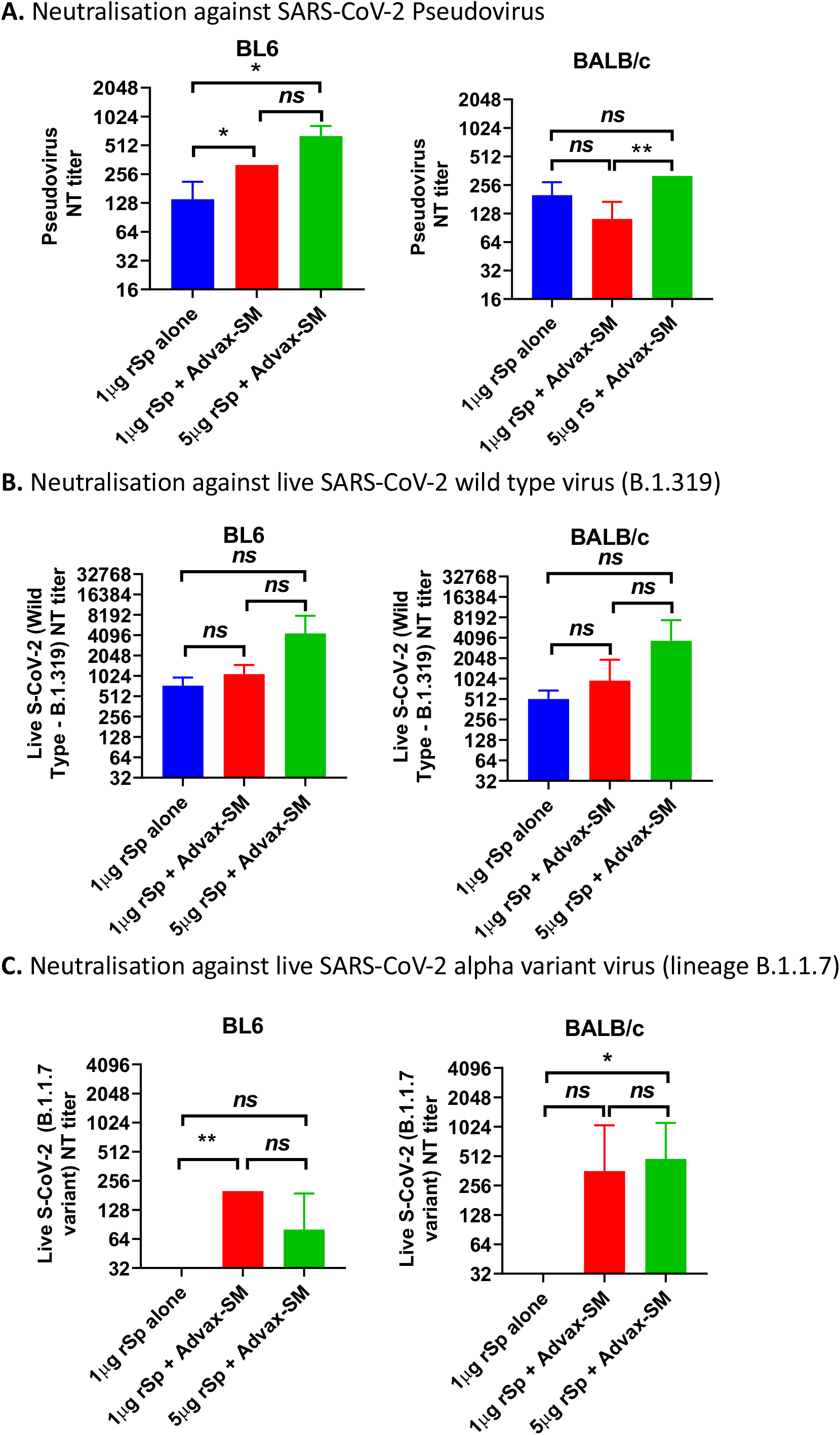
Neutralisation antibody titres from sera of immunised BL6 and BALB/c mice determined by (A) SARS-CoV-2 Spike pseudotyped lentivirus assay, (B) live SARS-CoV-2 wild type virus (lineage B.1.319) and (C) live SARS-CoV-2 variant of concern “Alpha” virus (lineage B.1.1.7, or “UK-strain”). Statistical analysis by Mann-Whitney test. *; p < 0.05, **; p < 0.01, ns; not significant.

Both BL6 and BALB/c mice immunised with Advax-SM adjuvanted rSp produced antibodies able to neutralise live SARS-CoV-2 virus. In BL6 mice, the highest neutralising antibodies were seen after immunisation with rSp 5μg+Advax-SM (GMT 3,712), then rSp 1 μg with Advax-SM (GMT 1088) and then rSp alone (GMT 736) (**Figure 3B**). The same trends were seen in BALB/c mice with highest response for rSp 5μg+Advax-SM (GMT 4,352), then rSp 1 μg with Advax-SM (GMT 960) and finally rSp alone (GMT 512). To evaluate potential cross-protection against a variant strain, sera were also tested for ability to neutralise live SARS-CoV-2 “Alpha” variant of concern (lineage B.1.1.7, or “UK-strain”). Only sera from BL6 or BALB/c mice immunised with Advax-SM adjuvanted rSp were able to neutralise the B.1.1.7 variant virus, with no neutralisation activity seen in mice immunised with rSp alone **(Figure 3C)**.

We next asked whether there was any correlation between total spike or RBD antibody levels and pseudotype or live virus neutralisation titres. In BL6 mice, there was a positive correlation between spike and RBD binding IgG and pseudotype and live virus neutralisation titres, with the highest correlation between spike IgG and pseudotype neutralisation (r^2^= 0.49, p<0.0035), followed by RBD IgG and pseudotype neutralisation (r^2^= 0.38, p<0.015) (**Supplementary Figure 3A**). Interestingly, there were only weak non-significant correlations between spike IgG and live virus neutralisation titres (r^2^= 0.20, p<0.09) or RBD IgG (r^2^= 0.22, p<0.07).

In BALB/c mice, there was a positive correlation for spike IgG with pseudotype neutralisation titres (r^2^= 0.46, p<0.005) (**Supplementary Figure 3B**). There was also a positive correlation for spike IgM with both pseudotype (r^2^= 0.42, p<0.009) and live virus (r^2^= 0.4, p<0.01) neutralisation. However, there was no correlation between RBD IgM and either pseudotype or live virus neutralisation, suggesting that IgM in BALB/c might neutralise SARS-CoV-2 through an RBD-independent mechanism.

Interestingly, there was only a weak correlation between pseudotype and live virus neutralisation (r^2^ = 0.17, p<0.02) **(Supplementary Figure 4)**. This could reflect that pseudotype neutralisation assays are unique at two levels, they utilise greater numbers of viral particles to enable cellular transduction and GFP expression, and only measure the consequence of a single round of spike-driven cellular fusion. By contrast, the live virus neutralisation assay measures inhibition of viral entry and productive infection over a 3-day period with repeated rounds of viral replication. Hence, each assay measures different but important parameters of viral infection, providing clues as to the ability of immune sera to neutralize first-round viral entry vs. a replicative infection

### Vaccine-induced T cell responses in mice

Cytokine production was measured in culture supernatants of rSp-stimulated splenocytes obtained from immunised mice. In BL6 mice, rSp-stimulated IL-2, IFN-γ and TNF-α was significantly higher in the Advax-SM group, consistent with their Th1 bias **(Figure 4A-C)**. Similarly, in BALB/c mice, there was higher rSp-stimulated IFN-γ and TNF-α in the Advax-SM group **(Figure 4A-C)**. In BALB/c mice, rSp-stimulated IL-4, IL-6 and IL-10 production was highest in the rSp-alone immunised group, which also exhibited low IFN-γ and TNF production, consistent with a Th2 bias **(Figure 4D-F)**. IL-17 was modestly increased in Advax-SM adjuvanted rSp groups in both BL6 and BALB/c mice **(Figure 4G)**. Overall, rSp-alone groups exhibited a Th2 cytokine bias, while Advax-SM groups exhibited a Th1 bias with an increased IFN-γ/IL-4 ratio **(Figure 4H)**. ELISPOT assays on splenocytes from immunised mice confirmed significantly higher frequencies of IL-2 and IFN-γ secreting T cells in response to rSp stimulation in the Advax-SM groups **(Figure 5A-B)**. Anti-spike IL-4-producing T cells were significantly higher in BALB/c mice, consistent with their Th2 bias **(Figure 5C)**. Anti-spike IL-17-producing T-cells were also higher in the Advax-SM group in BL6 mice **(Figure 5D)**.

**Figure 4:**
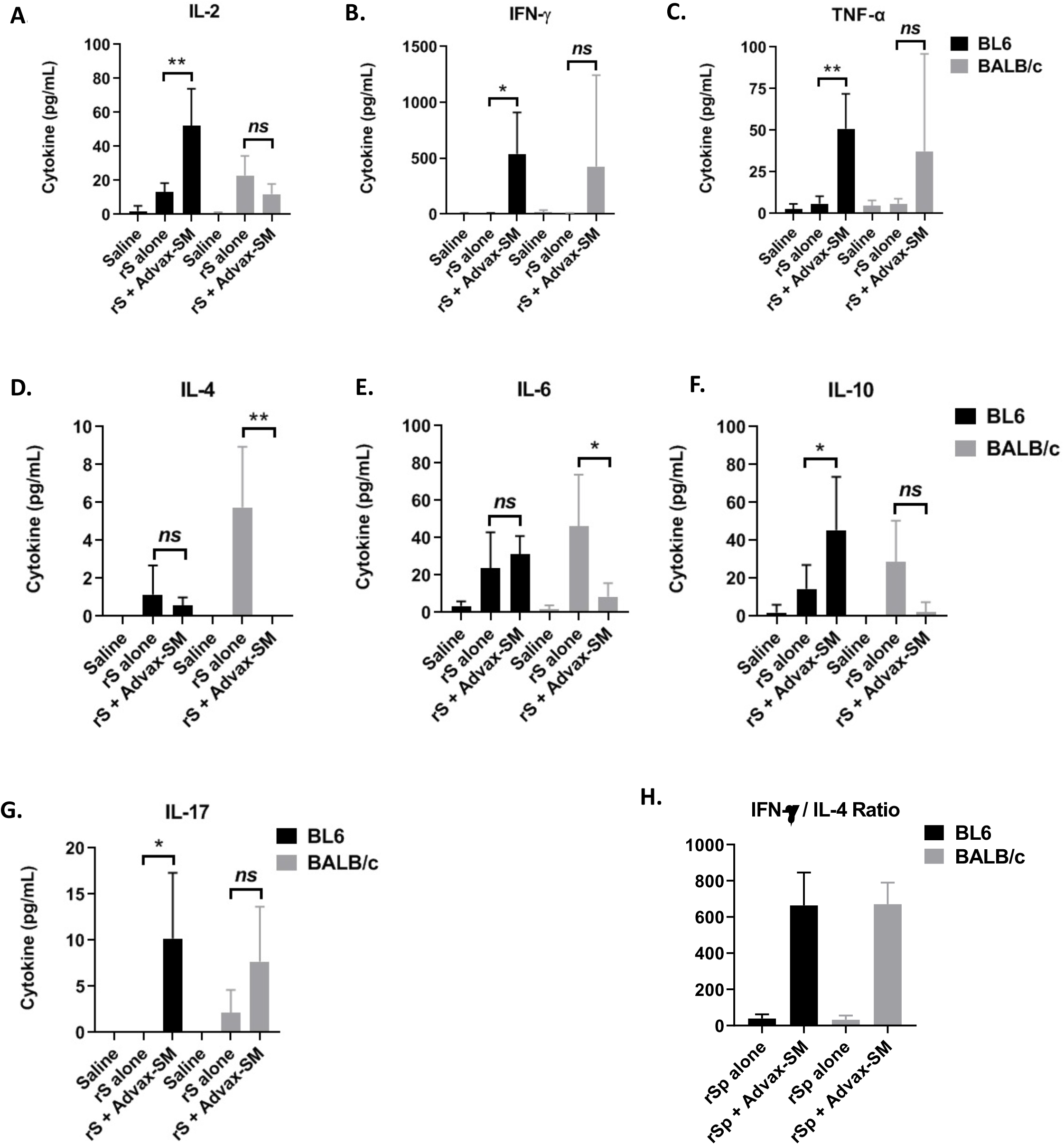
Co-administration of Advax-SM adjuvant enhances cytokine-producing cells in response to rSp antigen. Mice were immunised intramuscularly two times at 2 weeks apart with 1 μg rSp antigen alone or together with Advax-SM adjuvant. (A) Cytokines produced by splenocytes that have been collected 2 or 3 weeks after the last immunisation then cultured 2 days with rSp antigen were determined by CBA assay.

**Figure 5:**
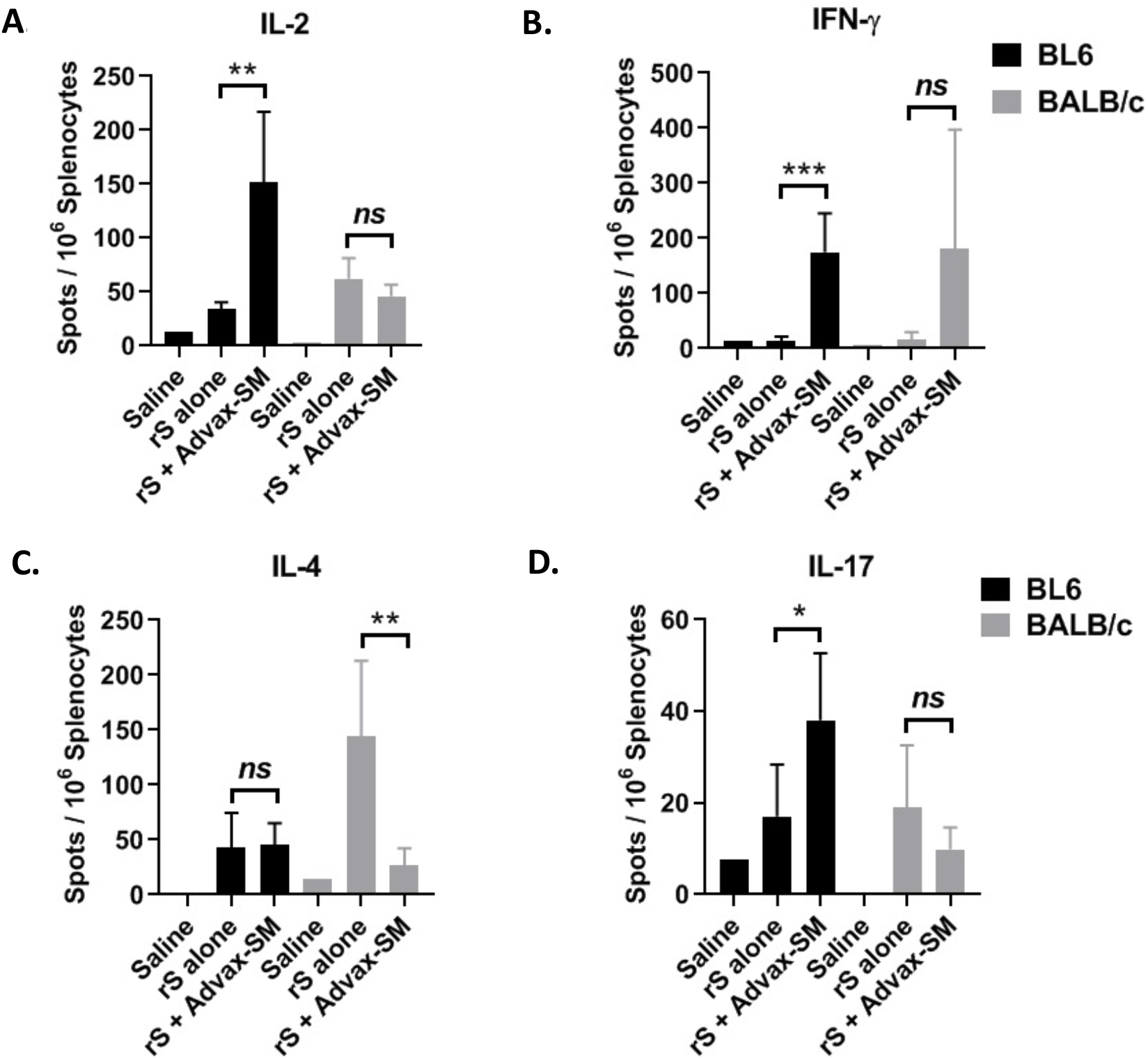
Antigen-specific cytokine-producing cells were evaluated following re-stimulation with rSp of splenocytes using anti-mouse cytokine antibody-coated plates by ELISPOT. Statistical analysis was done by Mann-Whitney test. *; p < 0.05, **; p < 0.01, ***; p < 0.001, ns; not significant.

Spike-specific CD4+ and CD8+ T cell memory cell population were further assessed using a CFSE-dye dilution proliferation in response to rSp stimulation. Notably, anti-spike CD8 T cell responses were markedly increased in both BALB/c and BL6 mice that had received Advax-SM adjuvanted rSp **(Figure 6)**, consistent with the high levels of anti-spike CTL activity also seen in mice receiving this formulation **(Supplementary Figure 1)**. There was also a clear trend to higher anti-spike CD4 T cell responses in mice that had received Advax-SM adjuvanted rSp, although this difference only reached significance in the Balb/c group **(Figure 6)**.

**Figure 6:**
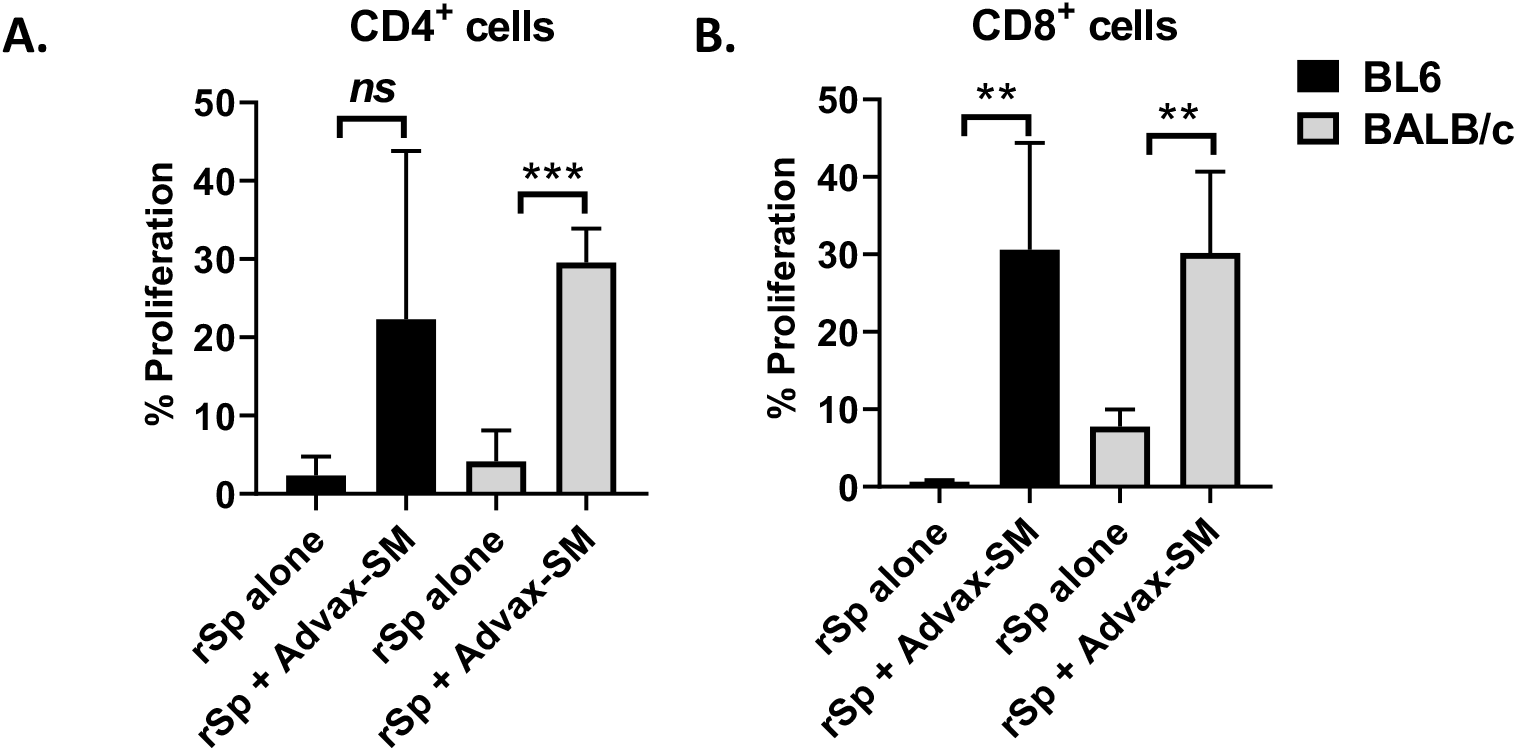
Antigen-specific proliferation was measured by FACS using CFSE-labelled splenocytes after culture for 5 days with rSp antigen. Results are presented as mean + SD. Statistical analysis was done by Mann-Whitney test. *; p < 0.05, **; p < 0.01, ***; p < 0.001, ns; not significant.

### Vaccine protection in ferrets

Having confirmed that the formulation of rSp with Advax-SM adjuvant gave optimal immunogenicity whether measured by neutralising antibody, T cell cytokines or CTL responses in mice, we next moved to test the efficacy of this optimised formulation in a ferret infection challenge model. Ferrets were given two immunisations 2 weeks apart with either of three dose levels of rSp protein (50, 25, or 12.5 μg) all formulated with the same dose of Advax-SM adjuvant, with control groups receiving two doses of an irrelevant influenza vaccine with the same dose of Advax-SM adjuvant (adjuvant control), two doses of saline (saline control) or just a single dose of rSp 50 μg + Advax-SM (single dose control) **(Figure 7A)**.

**Figure 7:**
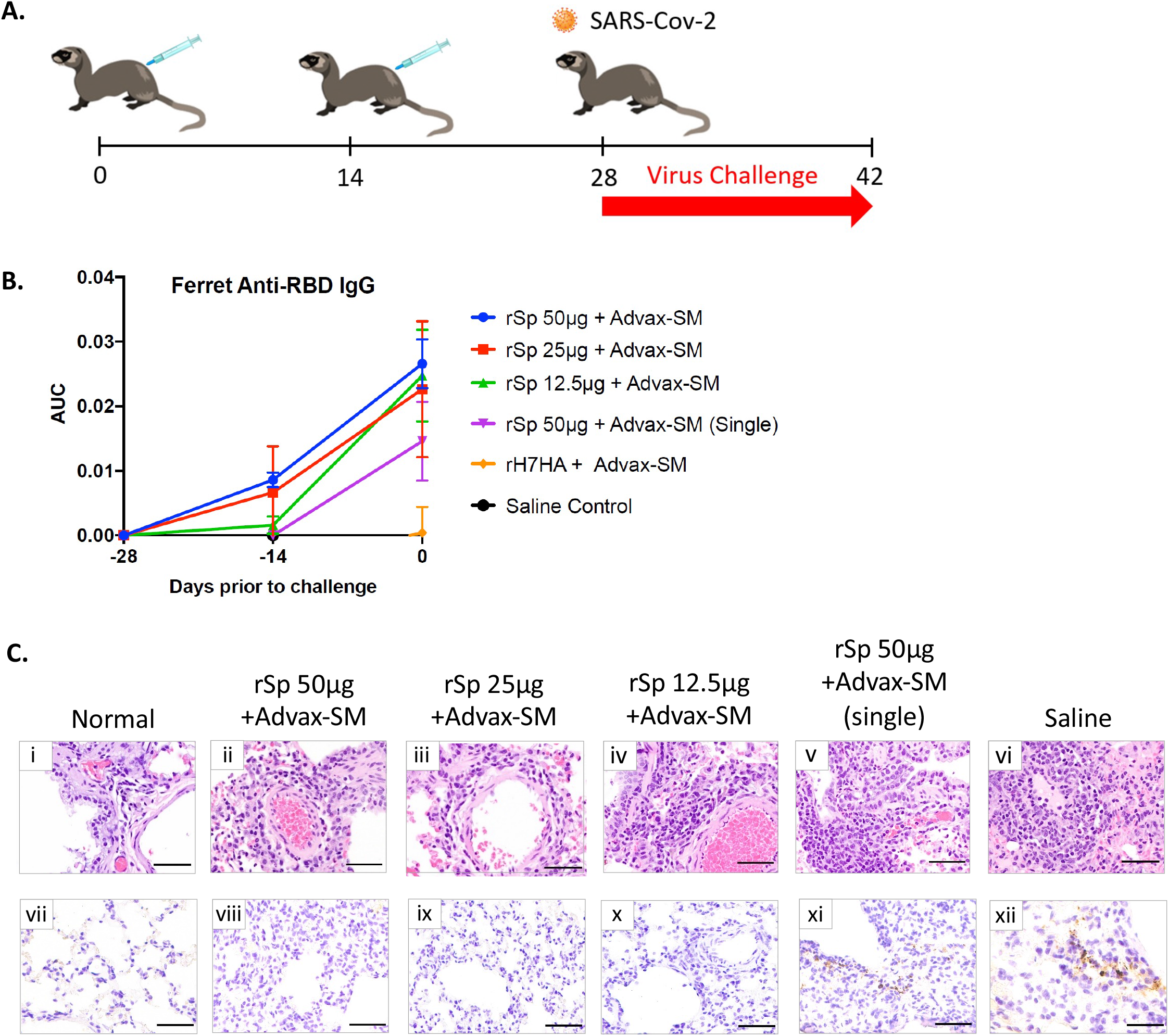
Ferret SARS-CoV-2 Challenge Model. (A) Overview of Ferret study design. Ferrets (n = 6) were vaccinated at day 0 and boosted at day 14 with SARS-CoV-2 spike protein (12.5, 25 or 50 μg) formulated with 15 mg Advax-SM adjuvant. Control ferrets either received saline only (n=3) or were immunised with influenza vaccine formulated with Advax-SM as a control (n=3). Ferrets were challenged with SAR-CoV-2 virus (1 × 10^5 PFU) 14 days after last immunisation. Blood was collected at day −28, −14. Lungs were collected 3 days post challenge for immunopathology. Lungs and nasal washes were collected 3 days post challenge for viral titres. (B) ELISA for Ferret anti-RBD IgG antibodies. (C) Haematoxylin & Eosin (H&E) (i-vi) and anti-SARS-CoV-2 nucleoprotein immunohistochemistry (IHC) staining (vii-xii) at 40x magnification (scale bar = 50μm).

Sera was obtained 2 weeks after the first and second immunisation and measured for IgG to RBD. Anti-RBD IgG was detectable even 2 weeks after the first dose in all rSp-immunised groups and levels were further increased 2 weeks after the second dose in those animals that received the second rSp dose **(Figure 7B)**. Control groups had negligible RBD titres at all time points. Two weeks after the second dose there was no significant effect of rSp dose seen on RBD titers between similar for the animals that had received wither the 50, 25, or 12.5 μg doses.

In response to the nasal virus challenge the ferrets showed minimal clinical signs of SARS-CoV-2 infection **(Supplementary Figure 5A)**. At 3 days post infection, lungs were harvested from three ferrets per group and scored for cellular infiltration and injury. Mock-vaccinated ferrets demonstrated a trend towards higher interstitial pneumonia based on H&E staining, and viral antigen was detectable in lung cells by immunohistochemistry **(Supplementary Figure 5B and Figure 7Cxii)**. Ferrets that received just a single vaccine dose similarly showed a trend toward higher interstitial pneumonia in the H&E staining and positive virus staining **(Supplementary Figure 5B and 7Cxi)**. By comparison, all groups that received two doses of rSp with Advax-SM had negative SARS-CoV-2 virus staining in the lungs consistent with protection **(Figure 7Cviii-x)**.

Next virus load was assessed in day 3 post-challenge nasal washes. Ferrets that received two immunisations of rSp at 25 μg or 50 μg with Advax-SM adjuvant had no detectable nasal virus as measured by TCID_20_ assay. Similarly, only 5 out of 6 (84%) of the ferrets that received two immunisations at the lowest 12.5 μg rSp dose had no detectable virus in their nasal washes, day 3 post-challenge. By contrast, 50% of ferrets in the control and single dose groups had detectable virus in their nasal washes, day 3 post-challenge.

## DISCUSSION

Vaccines normally take 10-15 years from discovery to final market approval [31]. To accelerate our COVID-19 vaccine development we made use of a well-validated protein manufacturing platform complemented by *in silico* modelling analyses. In this way, as soon as the SARS-CoV-2 genome sequence became available in Jan 2020 [20], we were able to identify the putative spike protein, model its structure and use docking programs to predict human ACE2 as the main receptor for the virus, as then confirmed by others [24]. This facilitated our rapid design of a recombinant spike protein antigen able to be produced as a soluble secreted protein in insect cells to which Advax-SM adjuvant was then added to produce the final vaccine formulation, which we named Covax-19.

We first evaluated the immunogenicity of the vaccine in BL6 and BALB/c mice, confirming Covax-19 vaccine was effective in inducing IgG and IgM antibodies against the spike protein with potent virus neutralisation activity whether measured by pseudotype or wildtype virus neutralisation assays. Notably sera from Covax-19 immunised mice were able to cross-neutralise the B.1.1.7 virus variant. In mice, the Advax-SM adjuvated rSp vaccine induced a strong Th1 response characterised by a switch from IgG1 to IgG2 and IgG3 IgG isotypes together with an increased frequency of IL-2, IFN-γ and TNF-α secreting anti-spike T cells, and a high level of CTL killing of spike-labelled target cells, *in vivo*. By contrast, immunisation with rSp alone (or formulated with Alhydrogel adjuvant) induced a predominantly Th2 antibody and T cell response against spike protein, with lower levels of neutralising antibody against the wildtype virus and no neutralising activity against the B.1.1.7 virus variant and no *in vivo* CTL activity against spike-labelled targets. Overall, this demonstrated that our insect cell expressed spike ECD construct when formulated with Advax-SM adjuvant is an effective immunogen against SARS-CoV-2.

SARS-CoV-1 and SARS-CoV-2 viruses target interferon pathways [32, 33]. Hence, coronavirus vaccines should ideally prime a strong memory Th1 and interferon response with CD8+ T cells playing a critical role in detection and silencing of virus-infected cells [34]. Immunisation with Advax-SM adjuvanted rSp induced a high frequency of spike-specific memory CD4+ and CD8+ T cells, which were not seen in mice immunised with rSp alone. This suggests that the Advax-SM adjuvant was able to induce effective dendritic cell cross-presentation of spike protein to CD8 T cells, with CD8 T cell priming to exogenous antigens typically requiring activation of CD8+ dendritic cells [35]. Notably, this CD8 T cell crosspresentation was associated with significant *in vivo* CTL activity against spike-labelled targets suggesting that our vaccine should be able to robustly control infection, not just through induction of neutralisating antibody but also through induction of CTLs able to efficiently identify and kill any residual virus-infected cells in the body. It has been difficult to identify non-reactogenic adjuvants that induce strong CD8+ T cell responses, making this a potential key advantage of Advax-SM when used in viral vaccines where strong CD8+ T cell responses are likely to be important to protection.

Whereas other adjuvant platforms might provide some nonspecific antiviral protection via activation of the innate immune system, this has not been a feature of the Advax-SM adjuvant. Notably, there was no suggestion of reduced disease in the challenged ferrets here that were injected with Advax-SM adjuvant plus an irrelevant influenza vaccine. Similarly, in a past SARS CoV vaccine study we did not see any nonspecific protection in mice injected with Advax-CpG alone [14], nor did we see nonspecific protection of Advax and CpG alone in a ferret studies of H5N1 influenza [36]. Hence, this data all supports the enhanced protection of Advax-SM adjuvanted vaccine being solely mediated by its ability to enhance the adaptive immune response to the co-administered antigen.

The role of RBD-binding and neutralising antibodies in SARS-CoV-2 protection remains unclear. Initially, there were concerns of the possibility of antibody-mediated disease enhancement (ADE), as seen in SARS, dengue, Respiratory Syncytial Virus and other viral diseases [37]. Reassuringly to date, there have been no reports of ADE in COVID-19 patients, although those with the most severe COVID-19 illness often have high RBD and neutralising antibodies [38], suggesting neutralising antibody may not be enough, by itself, and other mediators like CTLs may also be required to fully control SARS-CoV-2 infection. Furthermore, a spike protein vaccine using a large Human Immunodeficiency Virus-derived protein trimerization tag and formulated with MF59 squalene adjuvant was shown to induce serum neutralising antibody, but provided no protection against nasal virus replication in either the ferret or hamster challenge models [39]. This suggests, at a minimum, that serum neutralising antibody is not able to prevent nasal virus replication. Furthermore, convalescent plasma has not proved effective when administered to severely ill patients but instead can induce immune escape variants [40]. Hence antibodies by themselves may not be sufficient to prevent or reverse COVID-19 disease. In Phase 2 trials, LY-CoV555, a cocktail of two IgG1 antibodies appeared to accelerate the decline in viral load over time but ultimately did not demonstrate clinical benefit [41] with other experimental monoclonal treatments still undergoing human testing [42].

The gold standard for antibody assessment remains live wildtype virus neutralisation assays, as these directly measure the ability of antibody to block cellular infection. However, different cell types may be infected via different mechanisms, so use of different cell lines in these assays could still give varying results. VeroE6 is frequently used in virus neutralisation assays with viral entry in these cells primarily endosomal and driven through cathepsin cleavage of the spike protein. In contrast, entry of SARS-CoV-2 into nasopharyngeal cells is driven through TMPRSS2-mediated cleavage of spike [43]. Whilst primary ciliated or goblet cells from nasopharyngeal tissue might be the most physiologically relevant cell type to use in neutralisation assays, high-throughput serology screening using air-interface cultures is not feasible. Whilst pseudotyping assays can be performed outside of a BSL3 facility, they measure only a single round of spike-mediated cellular fusion and, hence, do not mimic a natural infection where there are multiple rounds of entry and replication. RBD-binding antibody assays work on the presumption that antibodies that block spike protein from binding to ACE2 should stop virus infectivity. However, in animal studies vaccines that have been shown to induce anti-RBD IgG titres have not prevented virus replication in the nasal mucosa, suggesting either that such antibodies fail to prevent virus binding or entry or that they fail to get access to the nasal epithelium. How do results compare for these assays? Previous studies on convalescent patients have reported a positive correlation between spike-specific IgG and both pseudotype virus and live virus neutralisation. In our study, there was a poor correlation between the pseudovirus and live virus assays, suggesting they measure different determinants of neutralisation. The live virus assay measures the ability to block cell infection by a small pool of viral particles across 3 days of culture. The pseudotype assay uses a large pool of virus particles as a surrogate for a single spike-driven fusion event. In our study, total spike antibody ELISA predicted pseudotype neutralisation better than the RBD-binding ELISA. Interestingly, in BALB/c but not BL6 mice, there was a positive correlation between total anti-rSp IgM (but not anti-RBD IgM) with neutralisation titres. Elite donors with high neutralisation titres in human convalescent cohorts surprisingly achieved this via anti-viral IgM (S. Turville, personal communication). There is still no established correlate of COVID-19 protection that has been confirmed in either animal models or humans. The fact that different assays seem to yield different results suggests that the identification of a correlate based upon simple antibody protection may not be straightforward.

Unless a vaccine is able to induce potent sterilising immunity, some SARS-CoV-2 virus will inevitably enter cells in the nasal mucosa, where antibodies will not be able to reach it and begin to replicate. In the face of uncertainty over antibody protection and rapidly waning circulating SARS-CoV-2 antibody levels, a strong CD8 T cell response with interferon production and CTL activity is likely to be important for virus control. A large body of clinical data demonstrates that reduced T-cell responses and production of Th1 cytokines, such as interferon and IL-2, are seen in patients with severe COVID-19 disease [44–47]. Moreover, the mode of action and protection of several SARS-CoV-2 vaccines has been linked to induction of type I interferon secretion by amplifying T cell memory formation and promoting B cell differentiation and survival [48]. Notably, our Covax-19 vaccine imparted a strong Th1 bias and robust T cell responses by virtue of the Advax-SM adjuvant. By contrast, alum and squalene emulsion adjuvants induce a strong Th-2 bias, which may not be as beneficial for COVID-19 virus control [49]. COVID-19 vaccine with alum adjuvant demonstrated a Th2-biased response with a low IFN-γ/IL-4 ratio [50], and wse similarly saw a strong Th2 bias of alum for spike protein in the current study. Notably, alum- and squalene-adjuvanted COVID-19 vaccines were both ineffective against nasal virus replication [39]. This contrasts strongly, with the ferret protection data shown here, where Advax-SM adjuvanted rSp, completely prevented both lung and nasal virus replication, an exciting finding as prevention of nasal virus replication could be the key to prevention of virus transmission. We are currently do not know the mechanism for the prevention of nasal virus replication in the ferrets by Advax-SM adjuvanted rSp, with the possibilities that is it due to CTL induced by the vaccine migrating to the nasal mucosa where they might then rapidly eradicate virus infected cells, or an ability of the adjuvant to induce neutralising antibodies with different functional properties that are better able to access the nasal environment and prevent infectivity of the virus, or both. Future studies will attempt to explore these mechanisms further.

A limitation of the current study was that ferrets do not exhibit weight loss or other signs of SARS-CoV-2 clinical infection [51], with no animal models fully reproducing the features of severe SARS-CoV-2 clinical infection in humans. Ongoing studies are testing our Covax-19 vaccine in other species including hamsters and non-human primates to see whether the effects of the vaccine on inhibition of nasal virus replication extends to other species. The current study also only assessed protection soon after immunisation and it will also be important to assess the durability of vaccine protection.

### Conclusion

The COVID-19 pandemic represents a significant evolving global health crisis. The key to overcoming the pandemic lies in the development of an effective vaccine against SARS-CoV-2 that ideally prevents transmission as well as serious disease. Recombinant protein-based approaches to COVID-19 offer benefits over other technologies including a 40-year record of safety and effectiveness including in very young infants, together with reliable large scale manufacture and high stability under typical refrigerated conditions [52]. By contrast, other available technologies, including nucleic acid and adenoviral vector platforms have a high level of reactogenicity and pose cold chain and other distribution challenges [53, 54]. This study showed that an Advax-SM adjuvanted rSp vaccine (Covax-19 vaccine) when administered as two sequential intramuscular doses several weeks apart induces strong anti-spike antibody and T cell responses in mice and was able to protect ferrets against SARS-CoV-2 replication in the lung and nose, with the possibility that prevention of nasal replication may signal an ability to prevent virus transmission. Future clinical trials will be required to assess how this promising animal data translates into human protection.

## Acknowledgements

The following reagent was deposited by the Centers for Diseases Control and Prevention and obtained through BEI Resources, NIAID, NIH: SARS-Related Coronavirus 2, Isolate USA-WA1/2020, NR-52281. The authors would like to thank the University of Georgia Animal Resources staff, technicians, and veterinarians for animal care. We also acknowledge the expert assistance of Johnson Fung and King Ho Leong with the endotoxin and CTL assays.

## SUPPLEMENTARY MATERIAL

**Supplementary Figure 1:**
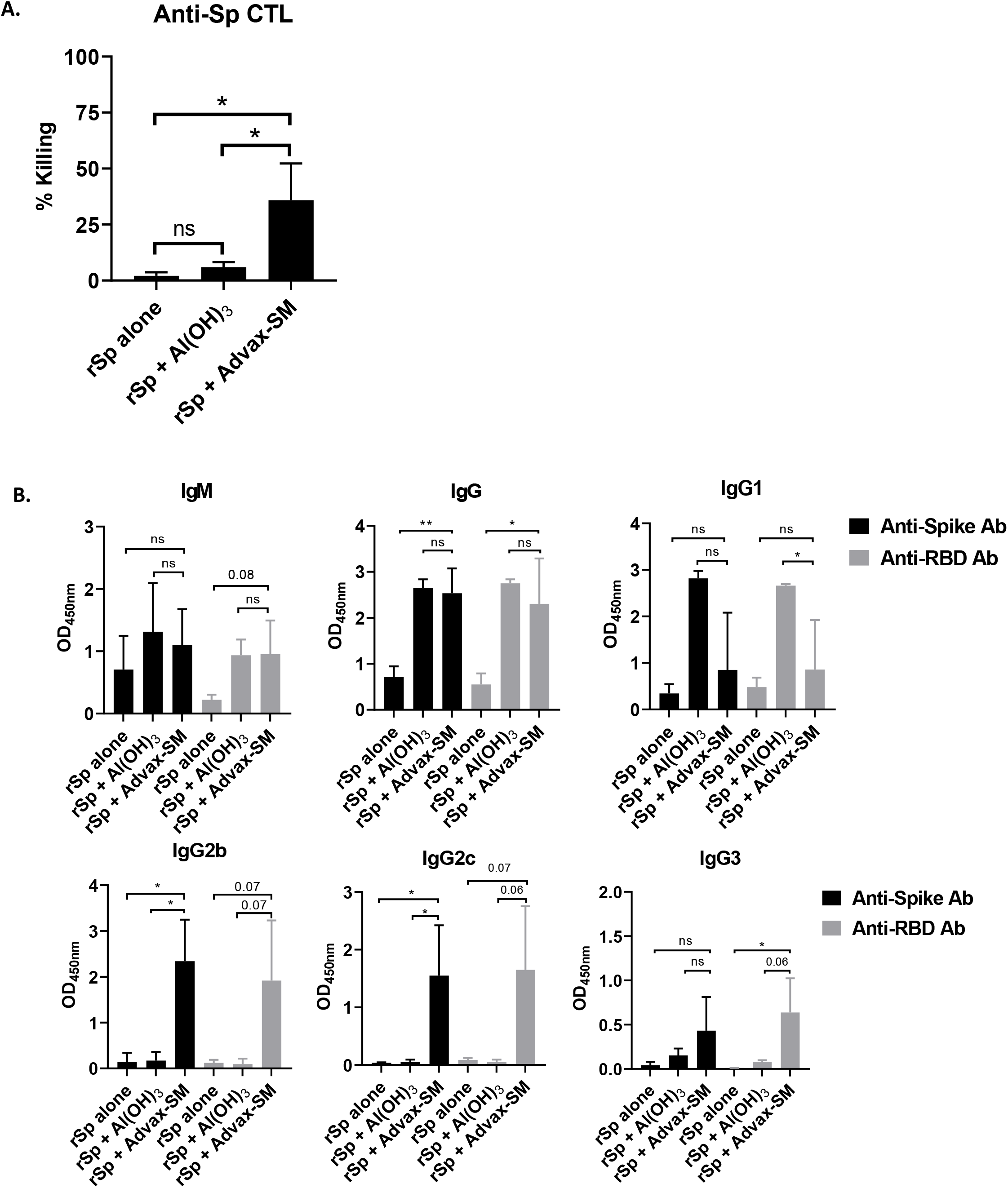
Advax-SM adjuvant enhances Cytotoxic T lymphocyte (CTL) activity and overcomes dominant Th2-bias compared to rSp antigen alone or together with Advax-SM adjuvant. Female BL6 mice were immunised i.m. twice at 2-week intervals with 1 μg rSp alone or with Advax-SM or 50 μg Al(OH)_3_ adjuvant. Blood samples were collected at 2 weeks and spleens at 4 weeks after the last immunisation. Shown are CTL functional assay (A) and ELISA (B) results (mean + SD). Statistical analysis was done by Mann-Whitney test. *; p < 0.05, ns; not significant.

**Supplementary Figure 2:**
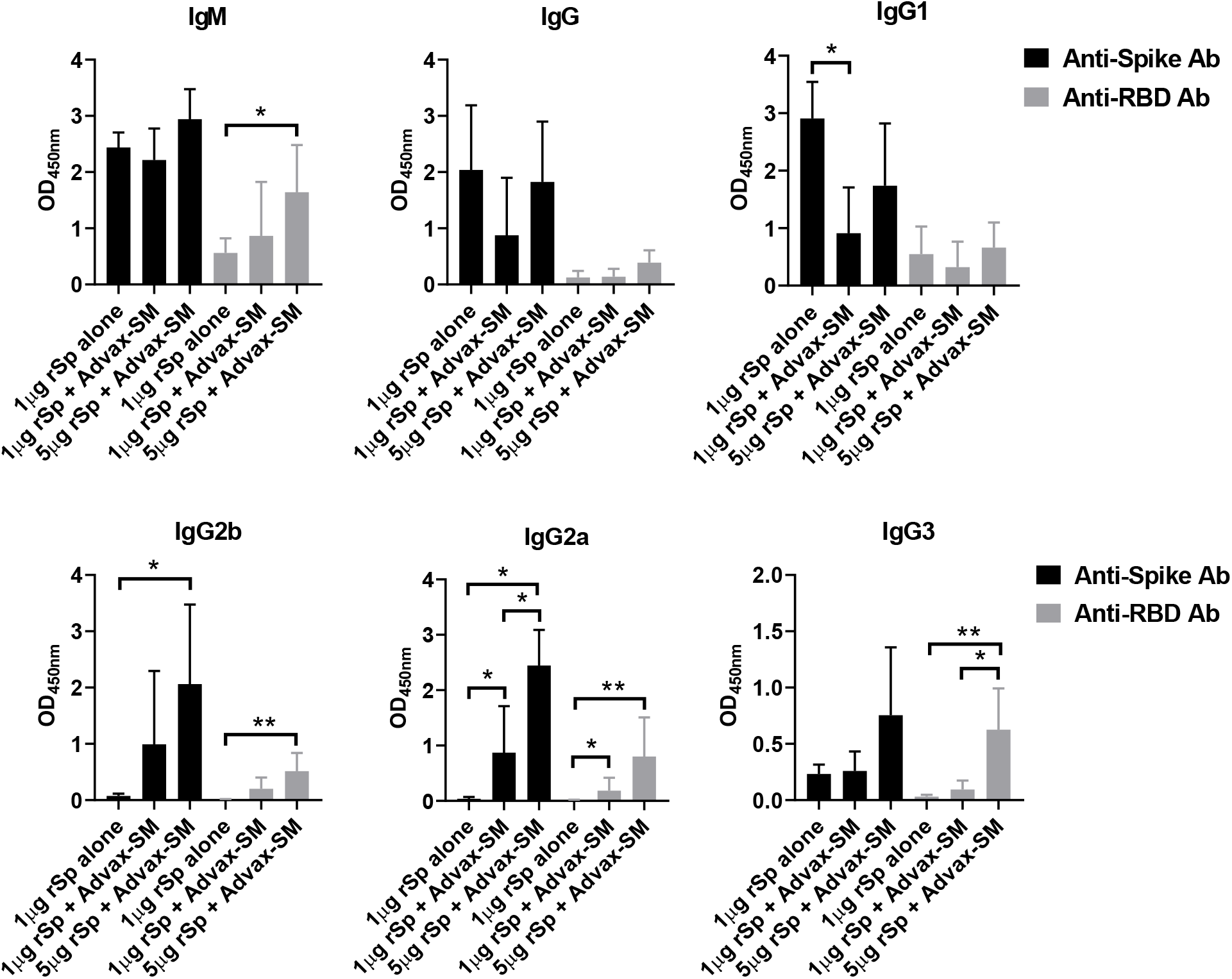
Co-administration of Advax-SM adjuvant enhances both vaccine-induced anti-rSp and anti-RBD antibodies in BALB/c mice. Shown are anti-rSp and anti-RBD antibodies measured by ELISA (mean + SD) in BALB/c mice. Statistical analysis was done by Mann-Whitney test. *; p < 0.05, **; p < 0.01.

**Supplementary Figure 3:**
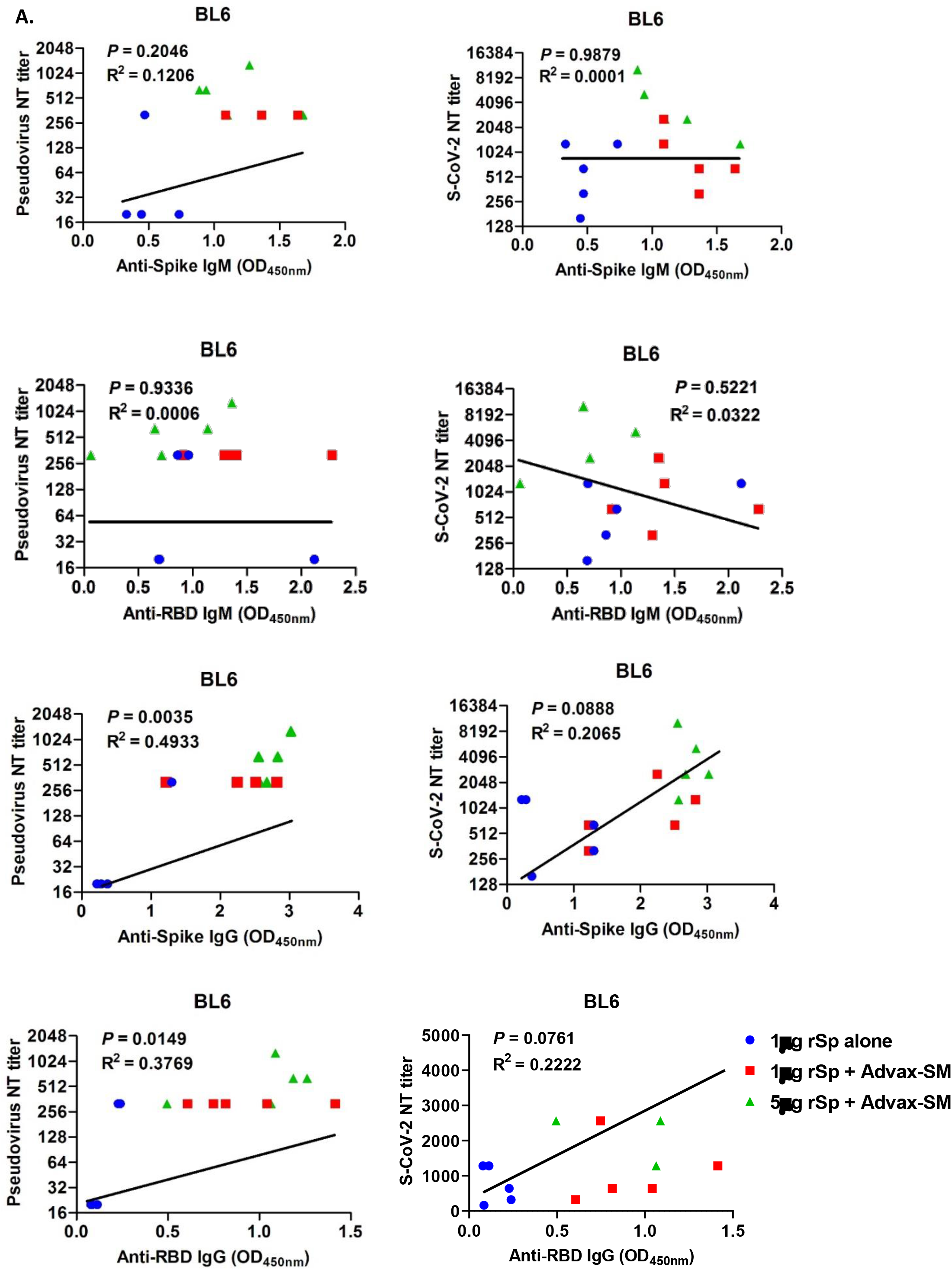
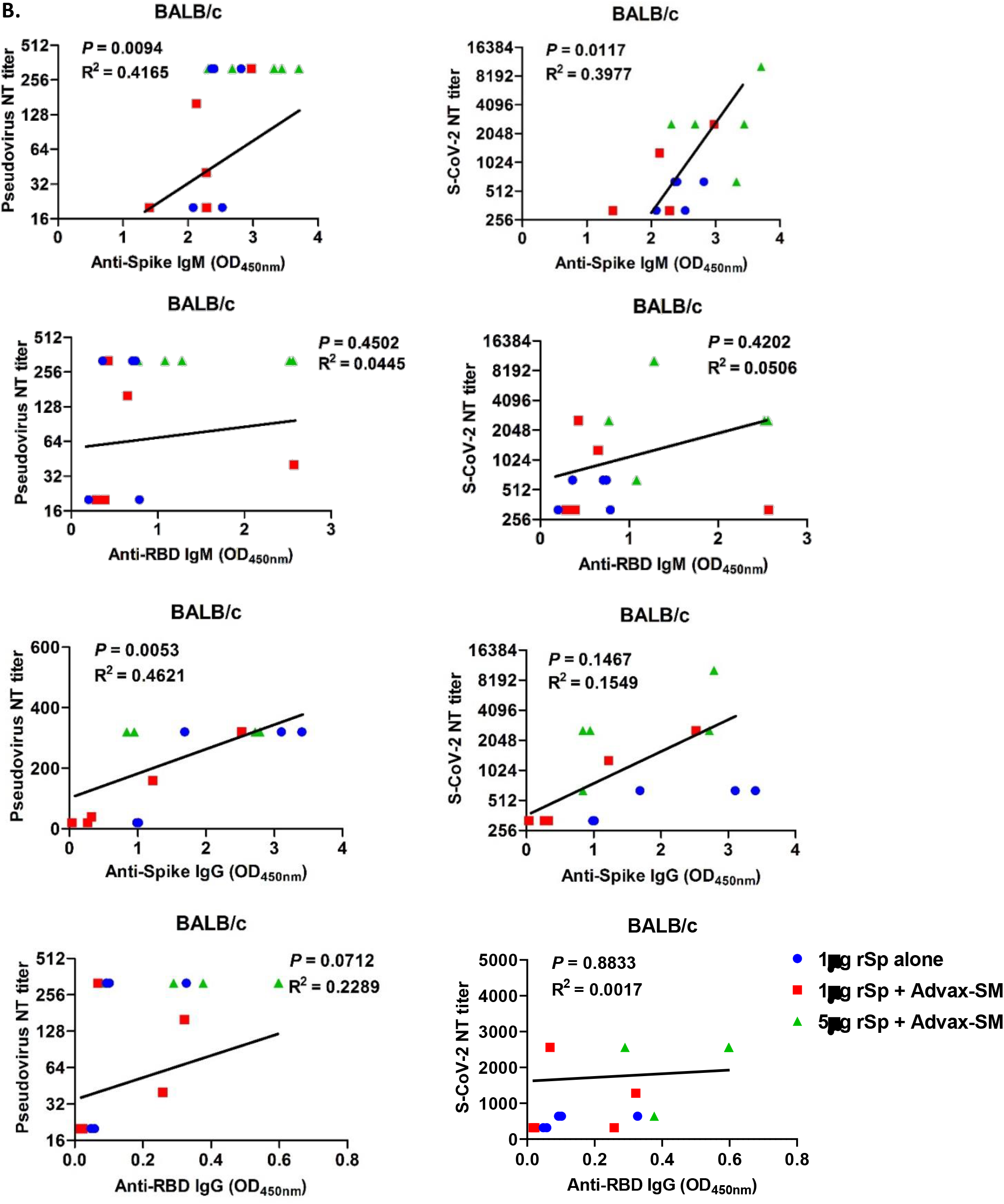
Neutralisation antibody titres determined by SARS-CoV-2 (S-CoV-2) Spike pseudotyped lentivirus-based platform and S-CoV-2 live virus. Correlation of NT titre and ELISA OD value in BL6 mice (A) and BALB/c mice (B).

**Supplementary Figure 4:**
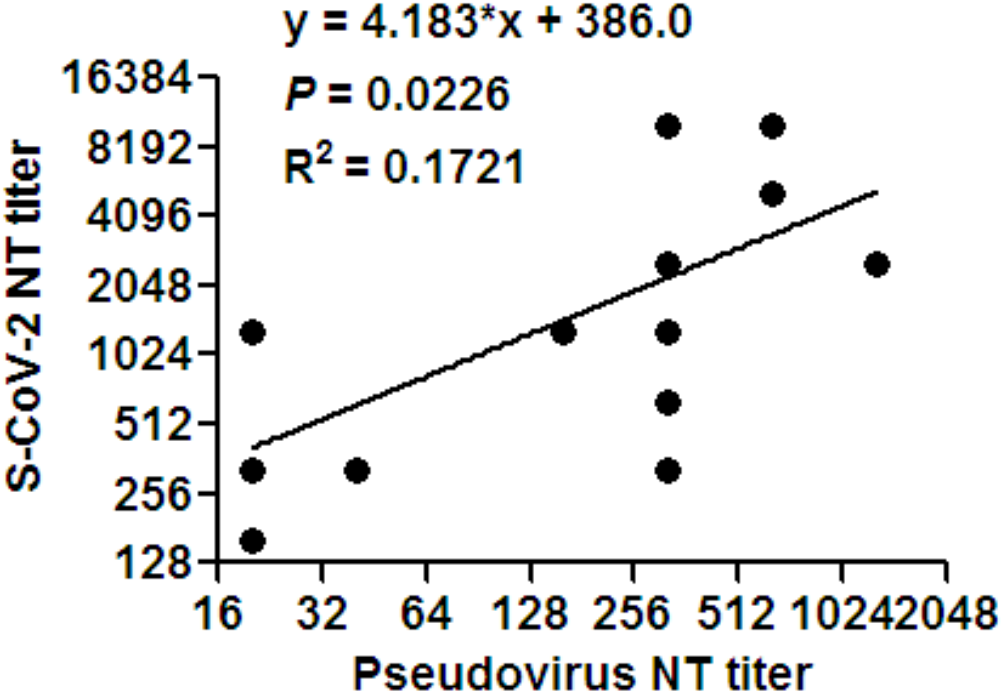
Correlation of S-CoV-2 and pseudotype neutralisation assays.

**Supplementary Figure 5:**
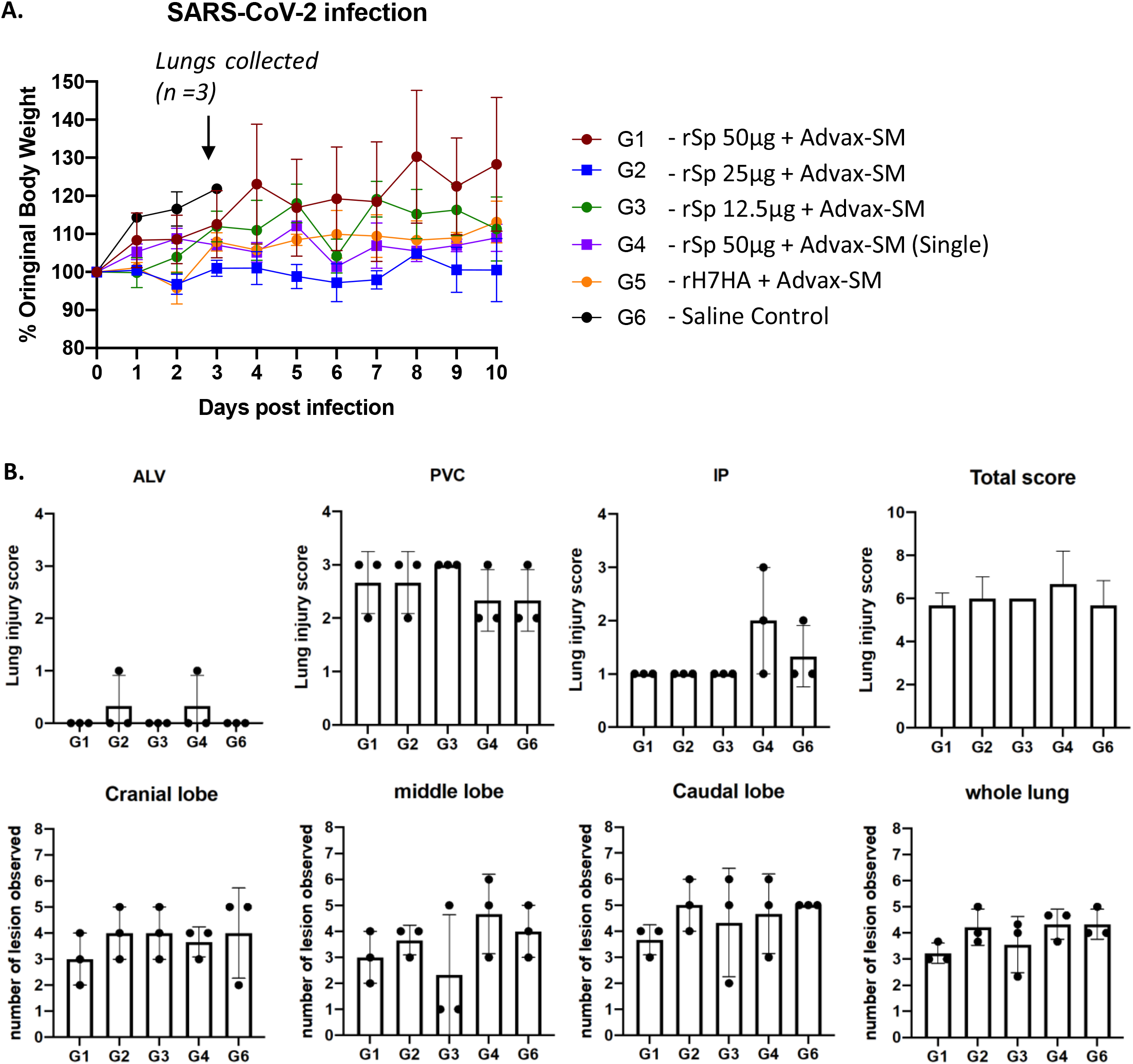
SARS-CoV-2 disease pathology in immunised and control ferrets postchallenge. (A) Change in body weight measured over 10 days post infection with 1 × 10^5^ PFU SARS-CoV-2 virus. (B) Lungs (n=3) were collected from each group at day 3 post-challenge, and histology sections were assessed by board-certified veterinary pathologist blinded to the groups for lung injury score and number of lesions.

